# Weak spatiotemporal response of prey to predation risk in a freely interacting system

**DOI:** 10.1101/215475

**Authors:** Jeremy J. Cusack, Michel T. Kohl, Matthew C. Metz, Tim Coulson, Daniel R. Stahler, Douglas W. Smith, Daniel R. MacNulty

**Author notes:** <u>Corresponding author</u> Jeremy J. Cusack – Biological and Environmental Sciences, University of Stirling, Stirling FK9 4LA, UK.

## Abstract

1. The extent to which prey space use actively minimises predation risk continues to ignite controversy. Methodological reasons that have hindered consensus include inconsistent measurements of predation risk, biased spatiotemporal scales at which responses are measured, and lack of robust null expectations.
2. We addressed all three challenges in a comprehensive analysis of the spatiotemporal responses of adult female elk (*Cervus elaphus*) to the risk of predation by grey wolves (*Canis lupus*) during winter in northern Yellowstone, USA.
3. We quantified spatial overlap between the winter home ranges of GPS-collared elk and three measures of predation risk: the intensity of wolf space use, the distribution of wolf-killed elk and vegetation openness. We also assessed whether elk varied their use of areas characterised by more or less predation risk across hours of the day, and estimated encounter rates between simultaneous elk and wolf pack trajectories. We determined whether observed values were significantly lower than expected if elk movements were random with reference to predation risk using a null model approach.
4. Although a small proportion of elk did show a tendency to minimise use of open vegetation at specific times of the day, overall we highlight a notable absence of spatiotemporal response by female elk to the risk of predation posed by wolves in northern Yellowstone.
5. Our results suggest that predator-prey interactions may not always result in strong spatiotemporal patterns of avoidance.

## Introduction

How, and to what extent, prey respond to the risk posed by predators are central questions in behavioural and community ecology (Sih, 1984; 1998). Although many types of behavioural responses, such as grouping (Hebblewhite & Pletscher, 2002; Fryxell et al., 2007) or increased vigilance (Elgar, 1989; Liley & Creel, 2007; Creel, Schuette & Christianson 2014; Creel et al., 2017, Dröge et al., 2017), can be studied through direction observation, others are more difficult to characterise unambiguously. In particular, the extent to which prey movement patterns actively minimise predation risk across space and time continues to ignite controversy (Creel et al., 2008). Indeed, there is a debate regarding the relative importance of proactive versus reactive spatiotemporal responses by prey to predators and the risk of predation (Creel, 2018). Proactive avoidance, where prey purposefully avoid areas or reduce activity during times of the day in which they are more vulnerable to predation (Prugh & Golden, 2014; Kohl et al., 2018), has been highlighted to a varying degree in a number of systems (Heithaus & Dill 2002; Creel et al., 2005; Fortin et al., 2005; Dupuch et al., 2009; Heithaus et al., 2009; Valeix et al., 2009; Padié et al., 2015). In contrast, reactive responses, which involve sudden displacements following more rapid changes in predation risk within the immediate surroundings have received increased attention in recent years owing to advances in tracking technology (Courbin et al., 2013; Middleton et al., 2013a; Basille et al., 2015; Courbin et al., 2016; Martin & Owen-Smith 2016).

Three common challenges arise when attempting to characterise prey spatiotemporal responses to predation risk. The first relates to how exactly predation risk is measured (Moll et al., 2017). It has often been assumed that the spatial distribution of a predator reflects a heterogeneous landscape of predation risk (Lima & Dill, 1990; Searle, Stokes & Gordon, 2008; Thaker et al., 2011). However, past studies have suggested prey may in fact be more likely to avoid specific habitats or landscape features that increase their vulnerability to predation (Hopcraft, Sinclair & Packer, 2005; Kauffman et al., 2007; Kohl et al., 2018). Predation risk may also vary over time, such as increase during times of the day when predators are more active or have higher hunting success rates (Palmer et al., 2017; Gehr et al., 2018; Kohl et al., 2018). In this context, Moll et al. (2017) recently recommended the use of multiple metrics in studies of predation risk.

A second complication lies in defining the spatial and/or temporal scale at which fear may act on prey behaviour (Kittle et al., 2008). A useful framework within which to consider this question was provided by Johnson (1980) in the form of a hierarchical classification of resource selection orders (see also Boyce, 2006). Past research investigating predator-prey interactions have primarily focused on whether the avoidance of predation risk by prey occurs at the level of home range selection (2^nd^ order) or at the level of patches within individual home ranges (3^rd^ order) (e.g. Courbin et al. 2013). However, few studies have considered how selection across these orders varies along a temporal dimension, for example 2^nd^ order selection between years or 3^rd^ order selection between different times of the day (although see Kohl et al., 2018).

A final challenge concerns how the expectation of behaviour in the absence of proactive and/or reactive responses is defined. For example, how would prey move through a given landscape if they ignored predation risk? Indeed, characterisation of prey spatiotemporal responses to predation risk has often been hindered by lack of an appropriate null model with which to generate expected behaviour, such as random movement (Gotelli & Graves 1996; Richard et al., 2013; Miller, 2015). Although step selection functions, which implement randomisations at the individual step level, provide a powerful tool to address this issue (Thurfjell, Ciuti & Boyce, 2014), their ability to randomise at the level of entire home ranges or to incorporate the temporal dimensions of space use is currently limited (although see Cozzi et al., 2018). An alternative method was recently proposed by Richard et al. (2013), who extended the application of null models used in community ecology to examine the potential for spatial interactions. They did this by randomly permuting and shifting roe deer *Capreolus capreolus* trajectories to obtain “pseudo-trajectories”, re-calculating the level of overlap with the distribution of female red deer (*Cervus elaphus*) to generate an expected distribution. Though promising, this approach has so far never been used to measure the strength of prey responses to predation risk.

In this study, we address all three challenges in a uniquely comprehensive analysis of the spatiotemporal responses of adult female elk (Cervus elaphus) to the risk of predation by grey wolves (Canis lupus) during winter in northern Yellowstone, USA. Since the reintroduction of wolves to Yellowstone in 1995, numerous studies have sought to characterise potential proactive versus reactive responses of elk and how these might relate to the trophic cascade observed across the ecosystem (Ripple & Beschta 2012). The majority of studies investigating movement and habitat selection responses by elk to the risk posed by wolves have revealed weak and inconsistent patterns (Fortin et al., 2005; Mao et al., 2005; Forester et al., 2007; Proffitt et al., 2009; White et al., 2008; Middleton et al., 2013a; Kohl et al., 2018). Despite this large body of research, which was drawn from multiple elk populations and relied primarily on movement data collected in the early years following wolf reintroduction, there remains a persistent contention that wolves have strong and consistent effects on elk space use (Winnie & Creel, 2017; Beschta, Painter & Ripple, 2018; Creel et al., 2018; Painter et al., 2018).

In this context, we first quantify spatial overlap between the winter home ranges of GPS-collared elk and three measures of predation risk: the intensity of wolf space use, the distribution of wolf-killed elk and vegetation openness. We then assess whether elk vary their use of areas characterised by more or less predation risk across the hours of the day. Lastly, we estimate encounter rates between collared elk and wolf packs during six 32-day winter periods occurring between 2013 and 2015. For all of these measures, we determine whether observed values are significantly lower than expected if elk movements were random with reference to predation risk. To do this, we implement a set of null model formulations that represent expectations of prey movement in the absence of predation risk effects, while accounting for elevation constraints known to affect winter elk movements. Using this approach, we answer the following questions:

1. Does elk choice of home range within northern Yellowstone (hereafter, philopatric behaviour) reflect proactive avoidance of spatial predation risk?
2. Does elk winter home range configuration reflect proactive avoidance of spatial predation risk?
3. Do elk minimise use of risky areas at specific times of the day?
4. Do elk avoid close encounters with wolf packs?

## Materials and Methods

### Study area

The northern Yellowstone winter range encompasses roughly 1,520 km^2^ of mountainous terrain and open valleys, with elevation ranging from 1,500 to 3,210 m (Houston, 1982). The area defines the winter range of seasonally migrating elk (White, Proffitt & Lemke, 2012; Tallian et al., 2017), and is largely composed of shrub steppe, with patches of intermixed lodgepole pine (*Pinus contorta*), Douglas fir (*Pseudotsuga menziesii*), Engelmann spruce (*Picea engelmanni*), and aspen stands (Houston, 1982; Despain, 1990). We consider wolf and elk trajectories recorded over the entire northern Yellowstone winter range – that is including land within Yellowstone National Park (YNP) and north of the park boundary – and hereafter refer to this as the northern range (NR). Winter severity in the NR is highly variable but in general snowfall increases from west to east due to an elevation gradient that approximates the distribution of elk on winter range, hence the inclusion of elevation in null model formulations (see below). Snow cover generally lasts from late October to early May but has recently become more variable (Middleton et al., 2013b).

Elk abundance in the NR has declined ~70% between 1995 and 2015. Currently, elk abundance numbers around 6000 individuals. It is estimated that only ~1500 of these elk overwinter in the YNP portion of the NR (Tallian et al., 2017). The decline in NR elk abundance has been largely due to a reduction in elk numbers within the NR’s YNP section (White & Garrot, 2005; White et al., 2012; Tallian et al., 2017). Elk are the primary prey of wolves in the study area (Smith et al., 2004; Tallian et al., 2017).

### Elk winter space use

We estimated individual-level home ranges for GPS-collared female elk during four winters (2012-13, 2013-14, 2014-15 and 2015-16) (Fig. 1a). A winter was defined as the period between November 1^st^ of a given year and 30^th^ April of the next. Elk collars (Iridium TrackM 3D, Lotek Wireless Inc.) were first deployed in February 2011, with new additions and redeployments occurring each subsequent winter. Adult (> 1 year old) female elk were captured using helicopter net-gunning. Recorded data were uploaded via Iridium satellite every 4-12 fixes and subsequently downloaded from a dedicated webserver. To ensure accurate representation of elk winter space use, we excluded winter movement paths for which the average fix frequency was more than five hours and the time difference between the first and last relocation was less than four months.

**Figure 1.**
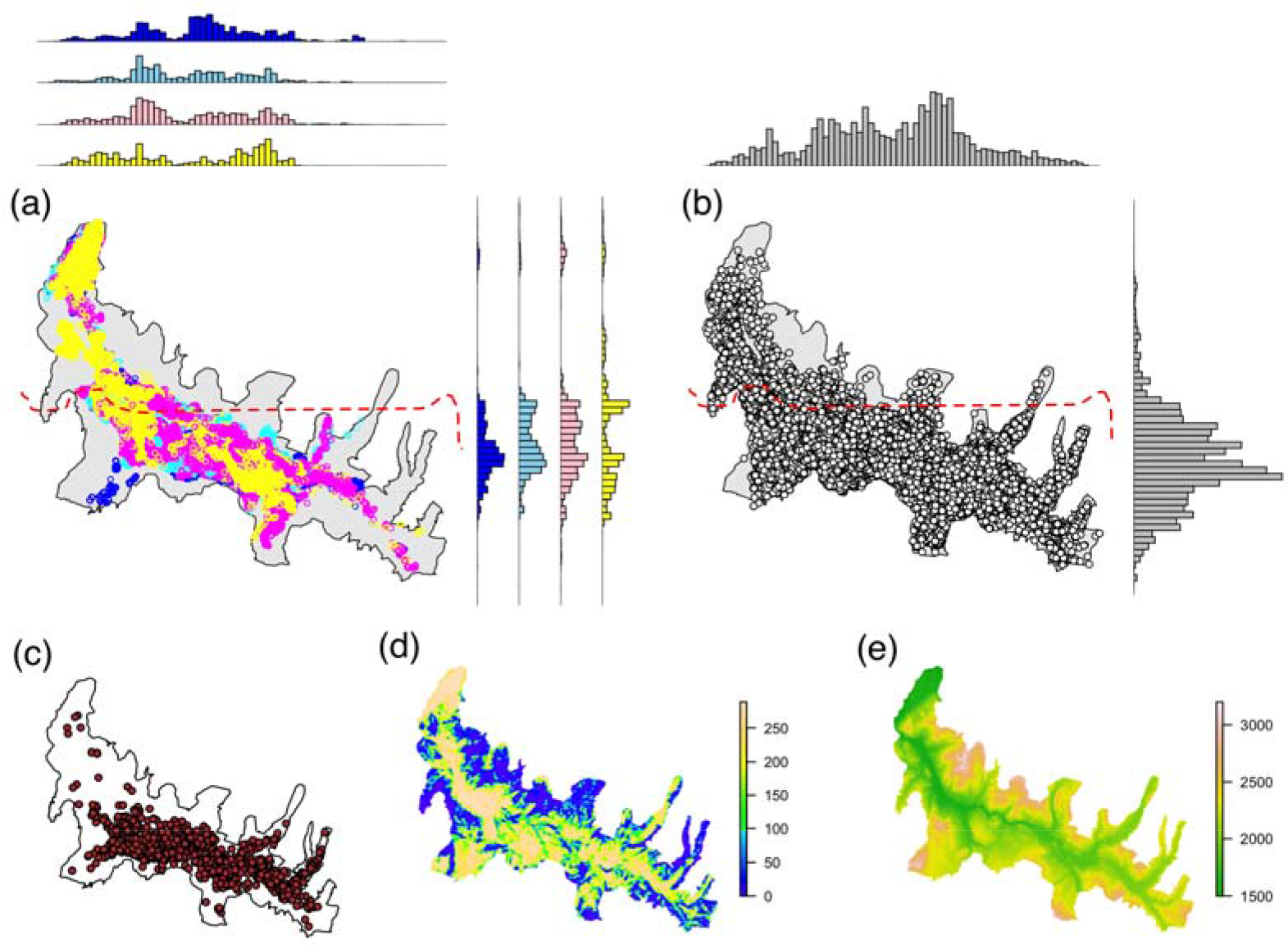
Overview of the spatial data collected across the Northern Range and used in this study. **(a)** Female elk GPS relocations for the winters of 2012 (yellow), 2013 (pink), 2014 (light blue) and 2015 (dark blue); **(b)** wolf GPS relocations recorded between 2004 and 2016; **(c)** distribution of elk female and calf kill sites; **(d)** vegetation openness (0=closed, 289=open); **(e)** elevation (in m). The dashed red line in (a) and (b) denotes the northern boundary of Yellowstone National Park.

For each winter, we estimated the individual-level utilisation distribution (UD) of each collared elk over a continuous grid of cell size 1 by 1 km using a Brownian bridge movement model (BBMM) implemented in the R package BBMM (Bullard, 1999; Horne et al., 2007). The BBMM is a continuous-time stochastic movement model, where the probability of being in an area is conditioned on 1) the distance and elapsed time between successive locations, 2) a measure of location error, and 3) an estimate of the animal’s mobility (the Brownian motion variance, see Horne et al., 2007). In other words, the model approximates the movement path between two subsequent locations by applying a conditional random walk. Because UD tails (i.e. beyond the 95 % isopleth) tend to be poorly estimated, we generated conditional 95 % UDs scaled to sum to unity (Benhamou et al., 2014). Location error for elk collars was unknown and fixed to a conservative estimate of 50 m. To avoid pseudo-replicating trajectories from collared elk belonging to the same group, we calculated an index of movement cohesion for every elk dyad within a given winter. We used Shirabe’s (2006) correlation coefficient, which measures the degree of correlation between the movement paths of two individuals as a multivariate Pearson product-moment correlation coefficient (Shirabe, 2006; Long et al., 2014). The index ranges from -1 (negative correlation) to 1 (positive correlation), with 0 indicating random movement. If two elk trajectories recorded during the same winter showed a movement correlation coefficient equal to or greater than 0.5, the one with the least number of relocations was excluded from the analysis.

### Wolf space use intensity

We used GPS collar data collected on wolves each winter between 2004 and 2016 to characterise long-term winter space use patterns by packs in the NR (Fig. 1b). Wolf GPS tracking has been routinely carried out by the Yellowstone Wolf Project since 2004, with a varying proportion of packs inside YNP sampled every year (details of collaring procedures can be found in Smith & Bangs, 2009). Although the exact model of fitted GPS collars varied during this period, all were manufactured by either Telonics (Mesa, AZ, USA), Televilt (Lindeserg, Sweden) or Lotek (Newmarket, ON, Canada). Average winter fix frequency between 2004 and 2016 varied between periods of intensive monitoring of wolf movements when relocations were obtained every hour (32-day winter periods, either Early Winter [EW] period between 14^th^ November and 15^th^ December or Late Winter [LW] period between 28^th^ February and 31^st^ March) and periods characterised by longer delays between relocations (average of 6 hours).

For each winter, we estimated a joint wolf UD representing the combined spatial activity of all collared wolves during that winter. The joint wolf UD was taken as the sum of individual wolf pack UDs – each of these weighted by the size of the corresponding pack (see Table S1, and Kauffman et al., 2007 for a similar procedure) – and scaled to sum to unity. Utilisation distributions were estimated using BBMMs estimated over the same spatial grid as that used for elk. We used a location error of 468 m for wolf packs as this represented the average distance between joint wolf movements. We assumed that this value accounted for the position of individuals that were not collared when estimating a pack’s UD (Benson et al., 2015). A final joint UD representing wolf long-term space use in the NR was then derived by averaging winter joint UDs and scaling to sum to unity. By averaging across winters – which differed in the number of packs collared (see Table S1 in Supporting Information) – we aimed to produce a space use pattern representative of where wolves were more or less likely to be encountered across the NR. Our study focuses on wolves collared south of the YNP boundary, and thus the estimation of the wolf UD in the northern section of the elk winter range relies on excursive movement from park packs.

### Elk kill site density and vegetation openness

We used a long-term, spatially explicit dataset on female elk and calf kill sites recorded in winter between 1995 and 2016 to derive a probability surface of observed predation by wolves (Fig. 1c). In a similar way to Kohl et al. (2018), we used a kernel density estimator implemented in the R package adehabitatHR to generate a smoothed spatial distribution of kill sites, setting a fixed bandwidth of 1000 m to match the resolution of the landscape grid. Lastly, we used a layer representing vegetation openness as a third measure of predation risk (Fig. 1d). Values in this layer ranged from 0 (thick forest) to 289 (open grassland) (see Kohl et al., 2018), which we subsequently standardised to sum to unity in order to ensure consistency with measures of wolf space use intensity and kill site density.

### Spatial overlap

We defined spatial overlap as the volume of intersection (VI) between the UD of a single elk during a given winter and a surface representing either one of the spatial predation risk indicators. We interpret VI as the proportion of the volume of the elk UD intersecting with a given predation risk layer (Kernohan, Gitzen & Millspaugh, 2001; Fieberg & Kochanny, 2005). The VI index, which ranges from 0 (no overlap) to 1 (complete overlap), has been widely used to compare UDs in a range of different taxa (Fieberg & Kochanny, 2005). In our case, if *UD_Elk_* and *UD_PR_* are the estimated utilisation distributions for an individual elk and predation risk type, respectively, then

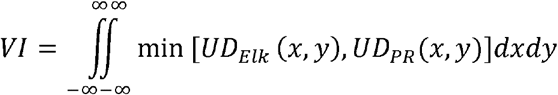

We calculated the VI index based on conditional 95 % UDs for elk, so as to minimise bias associated with the poorly estimated UD tails (Fieberg, 2007; Benhamou et al., 2014). We expected VI values to be low owing to the much larger spatial extent of predation risk layer values relative to that of individual elk UD values (i.e. there were many more instances of *UD_Elk_*(x,y) = 0 across the landscape, biasing VI towards 0). However, we stress that this in itself cannot be considered as evidence for proactive avoidance behaviour, and is the reason why we implemented a null model approach (see below).

### Hourly predation risk

To investigate whether elk use of risky areas varied across the 24-hour cycle, we modelled spatial predation risk level (wolf space use intensity, kill site density or vegetation openness) associated with a given relocation as a function of hour of the day. We used generalised additive mixed models (GAMMs) that included a term for first order auto-regressive processes (i.e. auto-correlation AR(1)), and implemented a cyclic cubic spline and Gaussian error structure (Wood, 2006). From this we obtained a prediction for the observed predation risk level associated with each hour of the day. For each type of predation risk considered, we ran one model per winter trajectory using the *gamm* function in the R package mgcv.

### Encounter rate

We measured the rate at which individual elk encountered wolf packs during six periods of intense monitoring (hereafter, winter periods) characterised by wolf relocations recorded every hour. Encounter rate was defined as *ST/n* where *ST* is the total number of recorded encounters with wolves and *n* represents the total number of fixes recorded for a given elk. Encounters consisted of spatially proximal and temporally simultaneous elk and wolf fixes defined according to specific distance *d* and time *t* thresholds, respectively (Long et al., 2014). We set *d* to 1000 m following Middleton et al. (2013a), who found that elk tended to increase their rates of movement, displacement and vigilance when wolves were within this distance threshold. Temporal proximity *t* was set to 1 hour as this represented the average length of a successful hunting bout by wolves (MacNulty, 2002). Thus, if elk and wolf relocations obtained in the same 1-hour window were observed to be within 1000 m of one another, they constituted an encounter. Importantly, we use the term “encounter” to denote a significantly increased likelihood of wolf-caused mortality (MacNulty, Mech & Smith, 2007), which we assume elk would actively avoid (Creel et al., 2005; Proffitt et al., 2009; Latombe, Fortin & Parrott, 2014). We excluded elk trajectories for which the number of tracking days was less than 30. For ease of interpretation, we present values of encounter rate for 100 elk fixes.

We modelled encounter rate as a function of the proportion of collared wolf packs in northern Yellowstone using a generalised linear mixed model (GLMM). The model response consisted of the number of encounters per trajectory with an offset term to account for varying number of fixes. We set the error distribution to Poisson and included elk ID as a random intercept to control for repeated measures on the same individuals across winter periods.

### Null model formulations

We used a null model approach to determine whether the observed spatial overlaps, encounter rates and hourly predation risk levels obtained for winter and period-level elk trajectories were less than expected by chance. All null model formulations were based on a correlated random walk, which randomly sampled the distributions of step lengths and turning angles derived from the observed elk trajectory to construct an alternative trajectory. We also imposed three constraints on null trajectories to ensure realistic outcomes. The first was that the generated trajectory fit within the same elevation range as the original trajectory (Fig. 1e). This was necessary to account for how deep snowpack excludes elk from high-elevation areas during winter irrespective of predation risk (Houston, 1982). Secondly, the null trajectory had to fit within the same bounding box area as the original. This ensured that the area covered by the trajectory did not affect expected outcomes. Lastly, null relocations could not occur outside of the NR.

To test whether philopatric behaviour by elk reflected avoidance of predation risk (Question 1), we generated null trajectories with starting locations sampled across the NR. Note that the starting location served as the centroid of the bounding box within which the null trajectory had to fit. We then constrained the starting location of null trajectories to a randomly sampled relocation from the observed trajectory, thus keeping the alternative elk trajectory within the same geographical area as the original (Questions 2 and 3). This latter formulation was also used to generate null trajectories for each winter period (Question 4).

For each winter and period-level elk trajectory, we generated 1,000 null trajectories, each time re-calculating the corresponding spatial overlap and encounter rate indices with each predation risk layer and period-level wolf trajectories, respectively. Hourly predation risk levels were re-calculated using the same null trajectories as for the spatial overlap analysis. Randomisations were carried out using the *NMs.randomCRW* function in the R package adehabitatLT (Calenge, 2006). Statistical testing consisted in computing the one-tailed probability *P* = (*k_e_* + 1)/*k* of getting a value equal to or less than the observed level, where *k* is the total number of null elk trajectories and *k_e_* is the number of values < observed. To control for the high number of significance tests, we applied a sequential Bonferroni correction by multiplying *P* by the number of elk trajectories in the corresponding winter, period or hour bin (Holm, 1979). We chose to implement a one-tailed test as we were interested in the alternative hypothesis of avoidance, which we refer to hereafter as a significant outcome. We report statistical significance at an α level of 0.05. All analyses were carried out in R version 3.5.0 (R Development Core Team, 2018).

## Results

### Spatial overlap

Elk winter UDs were estimated for 13, 22, 22 and 12 individuals during the winters of 2012, 2013, 2014 and 2015, respectively, totalling 69 winter trajectories. Trajectories showed a median of 181 days of tracking (range = 134 – 182) and an average of 2.39 hours between relocations (SD = 0.69) across all winters (Table 1, Figure S1). Movement correlation between contemporaneous trajectories was consistently <0.5. Wolf long-term space use across the northern Yellowstone elk winter range was estimated from 72,454 GPS relocations obtained from 23 individual packs (a total of 61 winter trajectories) between 2004 and 2016 (Table S1). A total of seven pairs of wolf trajectories exhibited a movement correlation coefficient greater than 0.5, resulting in the exclusion of the same number of trajectories prior to estimation of wolf space use intensity (Fig. 2a). The predation risk layer relating to elk kill site density (Fig. 2b) was derived from 1,780 wolf-killed elk detected between 1995 and 2016 across northern Yellowstone.

**Table 1.** Summary of winter elk trajectories.

| Winter | # trajectories | Total # relocations | Mean # tracking days | Mean # hours<br>between relocations (attempted interval) |
| --- | --- | --- | --- | --- |
| 2012-13 | 13 | 18,647 | 177.4 | 3.213 (2.5) |
| 2013-14 | 22 | 36,986 | 168.4 | 2.514 (2.5) |
| 2014-15 | 22 | 37,757 | 165.5 | 2.523 (2.5) |
| 2015-16 | 12 | 52,891 | 178.2 | 1.051 (1) |

**Figure 2.**
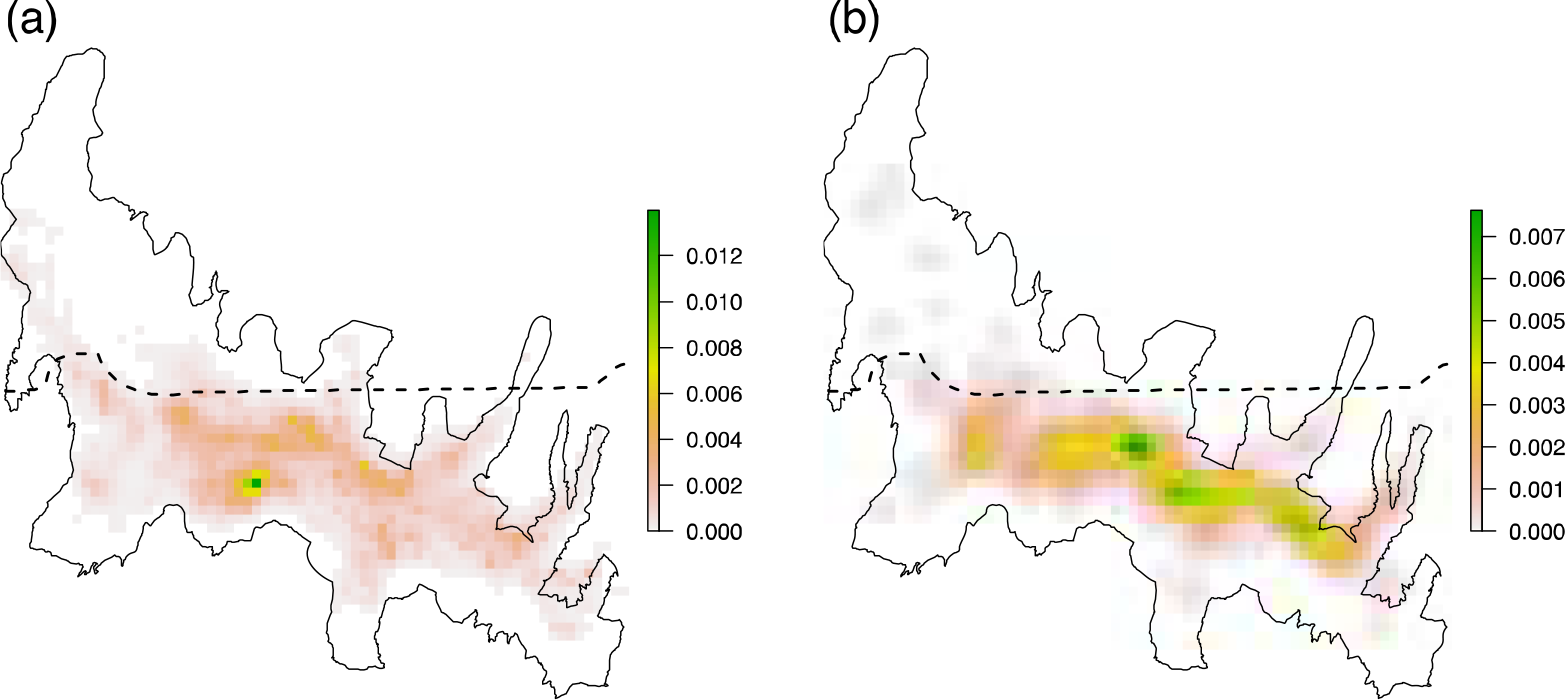
Predation risk layers representing wolf space use intensity **(a)** and elk kill site density **(b)**. The dashed line denotes the northern boundary of Yellowstone National Park.

As expected, spatial overlap values between elk winter home ranges and predation risk layers were low, ranging from 0.004 to 0.170 for wolf space use intensity, 0.007 to 0.361 for elk kill site density, and 0.006 to 0.058 for vegetation openness (see Tables S2 and S3). There was no evidence for proactive avoidance at the home range level when the null model formulation did not include a constraint representing philopatric behaviour, regardless of the predation risk layer. When philopatry was included in the null model formulation, 2 out of the 69 home ranges showed significantly less than expected overlap with vegetation openness, one in the winter of 2013 and the other in 2014. No home range displayed a significant outcome for wolf space use intensity or elk kill site density.

### Hourly predation risk

Across all hours of the 24-hour cycle, the mean percentage of individual elk using areas with lower than expected levels of predation risk was 1.4% (SD = 0.67) for wolf space use intensity, 0% (SD = 0) for kill site density, and 10.4% (SD = 2.4) for vegetation openness. For the latter metric, the proportion of significant outcomes was generally higher between 0700 and 1800, with a peak of 0.149 between 1200 and 1300 (Fig. 3).

**Figure 3.**
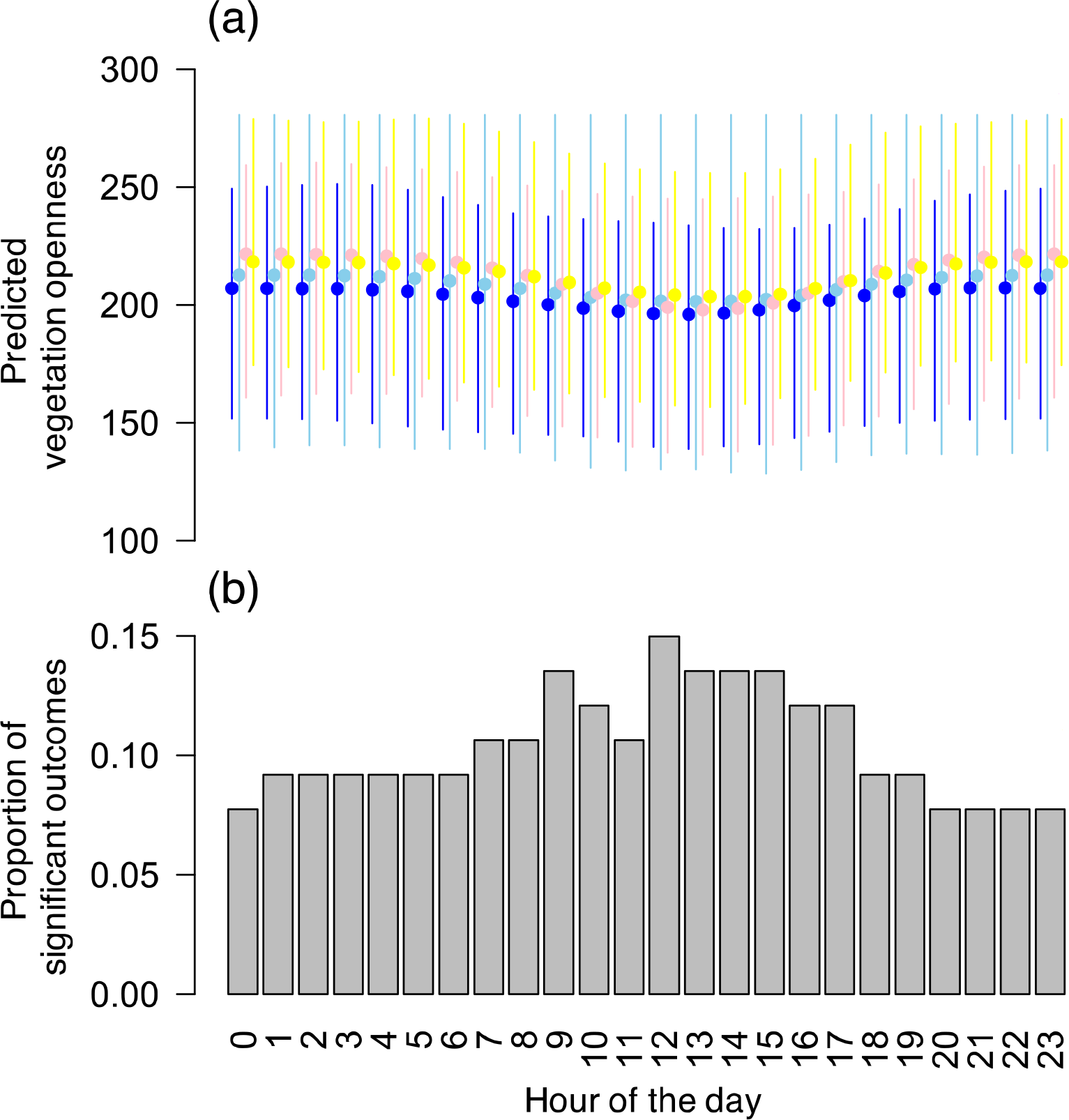
**(a)** Predicted mean level of vegetation openness per hour of the day. Full circles represent averages across individuals with bars showing the range of values. Colours indicate the different winters (yellow for 2012, pink for 2013, light blue for 2014 and dark blue for 2015). **(b)** Proportion of individual elk showing lower than expected mean vegetation openness per hour across all winters.

### Encounter rate

We recorded a total of 424 encounter events from 36,738 elk and 13,685 wolf pack relocations recorded across the six winter periods considered (Table 2). The majority of encounters (95.8%) were recorded inside YNP (Fig. 4a). For those elk that did experience encounters, these occurred on average once every 8.5 days with a range of 7.1 to 11.7 days across winter periods (Table 2). The shortest recorded distance between simultaneous wolf and elk relocations was 102.5 m. From the latter value, encounter frequency increased at a constant rate until the threshold of 1000 m (Fig. 4b). Encounters were more likely to be recorded during dawn (07:00 - 10:00) and dusk (16:00 - 19:00) than during the middle of the day or at night (Fig. 4c). Encounter rate increased significantly with the proportion of wolf packs collared within northern Yellowstone (GLMM; Fig. 4d and Table 2). Random intercept estimates showed a 12-fold variation across elk IDs, reflecting considerable differences in encounter rates at the individual level (Table S4). No elk trajectories were found to exhibit a lower than expected encounter rate with collared wolf packs. Note that a repeat of the analysis using a distance threshold of 500 m yielded the same result.

**Table 2.** Summary of winter period elk trajectories and encounter rates with collared wolves.

| Period<sup>*</sup> | # trajectories | Total #<br>relocations | Mean # hours<br>between<br>relocations | Total #<br>encounters | Encounter rate<sup>†</sup><br>median [range] | Mean # days<br>per encounter | # wolf packs collared<br>(proportion of active<sup>§</sup>) |
| --- | --- | --- | --- | --- | --- | --- | --- |
| LW 2013 | 18 | 5613 | 2.48 | 108 | 0.82 [0 – 5.56] | 8.4 | 2 (0.66) |
| EW 2013 | 18 | 5452 | 2.54 | 75 | 0.98 [0 – 4.22] | 9.2 | 3 (0.52) |
| LW 2014 | 15 | 4691 | 2.46 | 87 | 0.33 [0 – 3.91] | 11.7 | 3 (0.44) |
| EW 2014 | 24 | 7300 | 2.52 | 92 | 1.14 [0 – 6.17] | 7.1 | 4 (0.75) |
| LW 2015 | 23 | 7008 | 2.52 | 76 | 0.50 [0 – 4.56] | 9.1 | 4 (0.97) |
| EW 2015 | 22 | 6674 | 2.53 | 15 | 0.31 [0 – 1.05] | 8.3 | 2 (0.33) |

<sup>\*</sup>LW = Late Winter, EW = Early Winter
<sup>†</sup>Encounter rate per 100 fixes, i.e. the number of instances in which elk and wolf relocations within the same 1-hour window were within 1000 m of each other multiplied by 100
<sup>§</sup>Proportion of packs collared out of the ones known to be active in northern Yellowstone.

**Figure 4.**
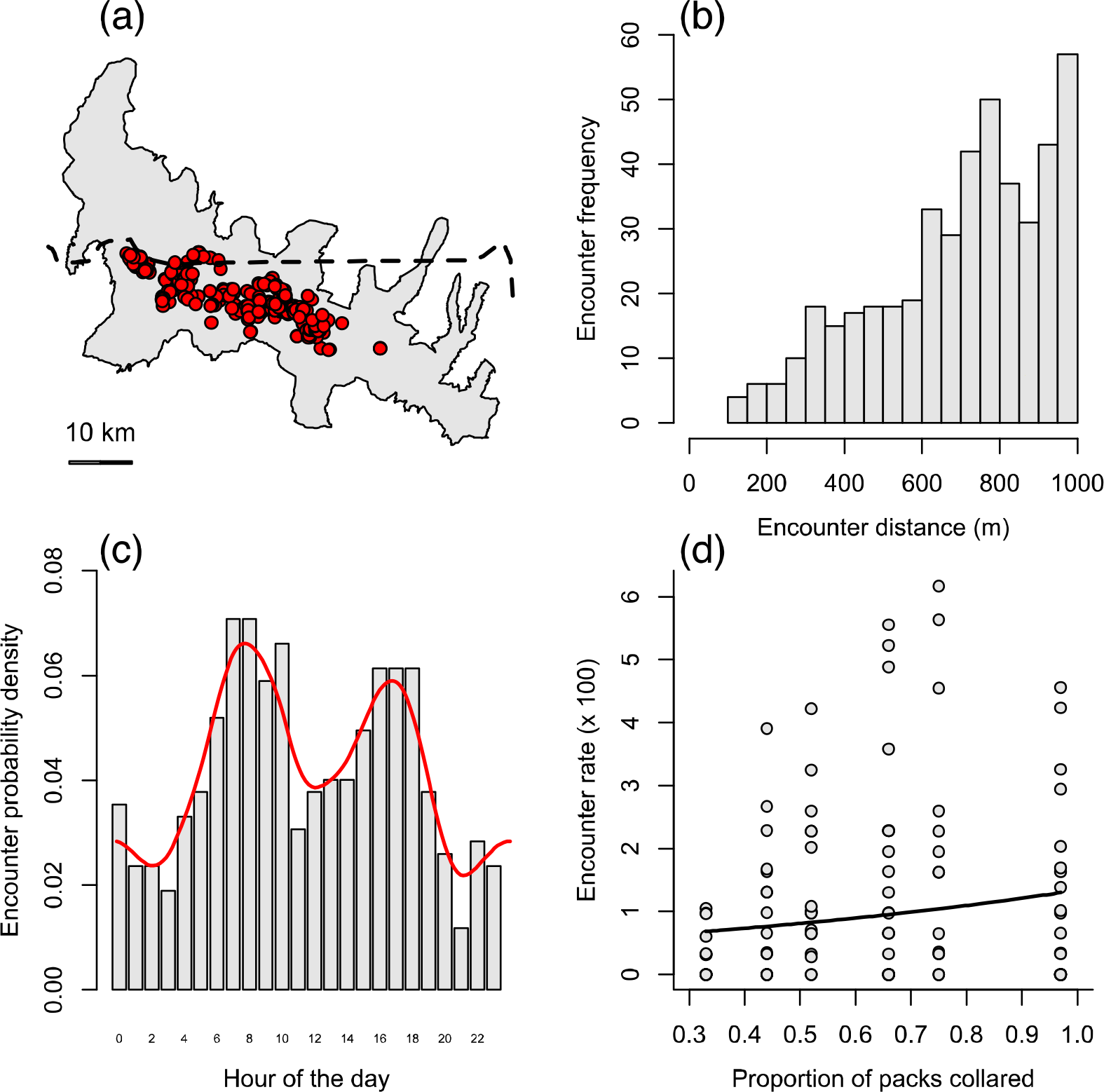
Details of encounter events recorded between GPS-collared elk and wolves in northern Yellowstone during six 32-day winter periods. These include the spatial distribution of recorded encounters (a), the frequency distribution of encounter distances (b), the probability density function of encounter times (c), and the relationship between encounter rate and the proportion of packs collared in northern Yellowstone (d). Encounters were defined as wolf and elk relocations obtained during the same 1-hour window and observed to be within 1000 m of one another. The dashed line in (a) denotes the northern boundary of Yellowstone National Park. The red curve in (c) represents the fitted density function. The fitted line in (d) was obtained from a Poisson generalized linear mixed model with the number of encounters as response variable, the proportion of collared wolves as explanatory variable, the number of fixes as an offset term, and elk ID as a random intercept. Encounter rate is expressed per 100 elk fixes.

## Discussion

Our study highlights a notable absence of spatiotemporal response by female elk to the risk of predation posed by wolves in northern Yellowstone. Home range selection by elk, both at the level of the entire NR and that defined by philopatric behaviour, did not reflect proactive avoidance of wolves themselves, nor of sites associated with a higher risk of being hunted successfully. Similarly, we found no evidence for reactive responses of individual elk to the presence of wolves in close proximity. Although a small proportion of elk did show a tendency to minimise use of open vegetation at specific times of the day (more so during the day than at night), in general we found a weak proactive temporal response to the different measures of predation risk. Together these results suggest that predator-prey interactions may not always result in strong spatiotemporal patterns of avoidance.

The limited proactive response of elk to wolf space use intensity concurs with findings from previous studies. In their comparison of elk movement patterns before and after wolf reintroduction, Mao et al. (2005) found that elk “did not spatially separate themselves from wolves” during winter months. One reason for this could be that elk are unlikely to be aware of the precise spatial distribution of a predator known to frequently course throughout their winter range (Bergman et al., 2006; Middleton et al., 2013a; Uboni et al., 2015). However, Kauffman et al. (2007) highlighted a discrepancy between kill site occurrence and wolf distribution, making the more general point that predator density may not be a good indicator of predation risk. To counter this criticism, we considered two additional measures of predation risk (Moll et al., 2017). These reflected the notion that elk might select for sites that reduce their vulnerability to being hunted successfully, such as areas of increased vegetation cover (Creel et al., 2005; Fortin et al., 2005). Yet, contrary to previous work, we did not find any evidence to support a proactive response to any of the predation risk measures, thus strengthening the idea that home range selection by elk in our study did not reflect avoidance of predation risk.

Recent work on the responses of prey to predators has highlighted the importance of time in modulating spatial relationships between prey movements and predation risk (Creel et al., 2008, Palmer et al., 2017). In particular, Kohl et al. (2018) recently revealed a dynamic landscape of fear, whereby elk use of risky areas in northern Yellowstone was dependent on wolf diel activity. Although the proportion of elk using open vegetation less than expected by chance did vary across the 24-hour cycle in the present study, this behaviour only concerned a very small proportion of the individuals tested each hour of the day. Interestingly, however, the detected avoidance response tended to be stronger during daylight hours, when wolves are more likely to be actively hunting (Vander Vennen et al., 2016; Kohl et al., 2018). The study by Kohl et al. (2018) used elk relocation data collected over the period 2001-2004 and it is possible that changes in elk behaviour towards wolves might have led to the weaker patterns observed in the present study. Elk numbers in northern Yellowstone were also much higher during the early years of wolf re-colonisation (MacNulty et al., 2016), when elk could have been using riskier habitats as a means to avoid safer but more crowded ones (i.e. density dependent effect).

The near absence of elk trajectories showing a lower than expected encounter rate with wolves is a surprising outcome of our study. From an ecological perspective, it is possible that other factors not considered here, such as elk group size (Gower et al., 2008; White et al., 2012), switches in habitat use (Creel et al., 2005; Fortin et al., 2005; Hernández & Laundré, 2005), and wolf pack size (MacNulty et al., 2012) allow individual elk to minimise predation risk despite close proximity to wolves, thus dampening small scale spatial avoidance patterns. Individual elk – and adult females in particular – might also tolerate close proximity to wolves because they frequently survive their encounters with them (MacNulty et al., 2007; MacNulty et al., 2012; Mech, Smith & MacNulty, 2015). From a methodological standpoint, we also have to consider the possibility that our definition of an encounter poorly described immediate predation risk, and that reactive avoidance occurs at a spatial scale < 500 m. Few high resolution relocation datasets are currently available that combine simultaneous predator-prey trajectories, and our study is valuable in developing a methodological framework within which these could be considered once they become more widely available.

Importantly, our findings are consistent with two key predictions of the predator-prey shell game occurring in a freely interacting system (Lima 1998; Mitchell & Lima, 2002). One of these relates to attempts by predators to get closer to prey. In a system such as northern Yellowstone where the winter movement of elk is constrained by philopatric behaviour and snow cover (Nelson et al. 2012), wolves may be better able to align their space use with that of their prey. A consequence of this would be the dampening of any potential avoidance patterns displayed by elk (as per Sih 1984; 2005), which might explain their overall absence in the present study. Another prediction states that prey should attempt to be unpredictable in space, and the lack of consistent movement patterns observed in the present study could be interpreted as a reflection of this. We emphasise that the methodology presented here, combined with other approaches such as step selection functions (e.g. Cozzi et al., 2018), could be used to assess behavioural responses on both sides of the predator-prey race.

We must acknowledge the potential limitations of our study. In particular, Creel, Winnie & Christianson (2013) recently reviewed sources of bias associated with the estimation of encounter rates between mobile predators and prey, some of which are relevant to the present study. Firstly, the fix frequency used to record elk movement trajectories, which averaged 2.39 hours across winters, may have led us to overlook instances of close proximity with wolves, and even entire hunting episodes (MacNulty, 2002; MacNulty et al., 2007). Although we cannot exclude this with absolute certainty, the 1-hour temporal window used to define encounters is likely to have minimised this problem. Secondly, not all of the packs active in the northern range during a given winter period were considered, which may have exacerbated the under-estimation of encounter rates. Nevertheless, our study considers movement trajectories from members of many of the dominant packs in northern YNP, and although the proportion of packs collared did positively influence observed encounter rate, it did not affect the absence of significant outcomes. Thirdly, we did not make use of more complex measures of dynamic interaction between simultaneous trajectories (reviewed by Long et al. 2014). Instead we chose to use a more intuitive measure of encounter rate, which we complemented with an assessment of significance based on values obtained under the assumption of random movement (Miller, 2015).

In summary, not only does our study provide a comprehensive assessment of the spatiotemporal response of individual prey to predation risk, but it also extends the use of null models to infer on interactive behaviour between different species. In doing so, it emphasises the challenges of detecting strong spatiotemporal responses by prey, and suggests that other factors relating to both predator and prey behaviour may be more important in shaping observed outcomes. Although our data were based on a system that has undergone extensive study over the past two decades, the considerations we highlight are particularly relevant to telemetry studies carried out in poorly known landscapes, in which spatial data are increasingly the first to be collected. In such cases, a clear understanding of species interactions, such as the proactive and reactive responses of prey to predators, may have to be gained through a combination of high-resolution GPS telemetry and direct observation.

## Acknowledgements

This work was supported by UK Natural Environment Research Council grant NE/J016527/1 to T. Coulson. Funding was also provided by the National Science Foundation (DEB-1245373), Yellowstone Park Foundation, the Tapeats Fund, the Perkin-Prothro Foundation, an anonymous donor, and the National Park Service. We thank Erin Stahler for assistance with the data collection and management, Chris Carbone, Marcus Rowcliffe and three anonymous reviewers for comments on previous versions of this manuscript.

## Authors’ contributions

JC designed the study; MK, MM, DRS, DWS and DM collected and shared the data; JC and TC performed the modelling work and analysed output data. JC wrote the manuscript, and all authors contributed substantially to revisions.

## Data accessibility

Data used in this study will be made available on a Dryad repository upon acceptance.

