## Supplementary figures and images for "Weak spatiotemporal response of prey to predation risk in a freely interacting system"

### Supplementary Materials

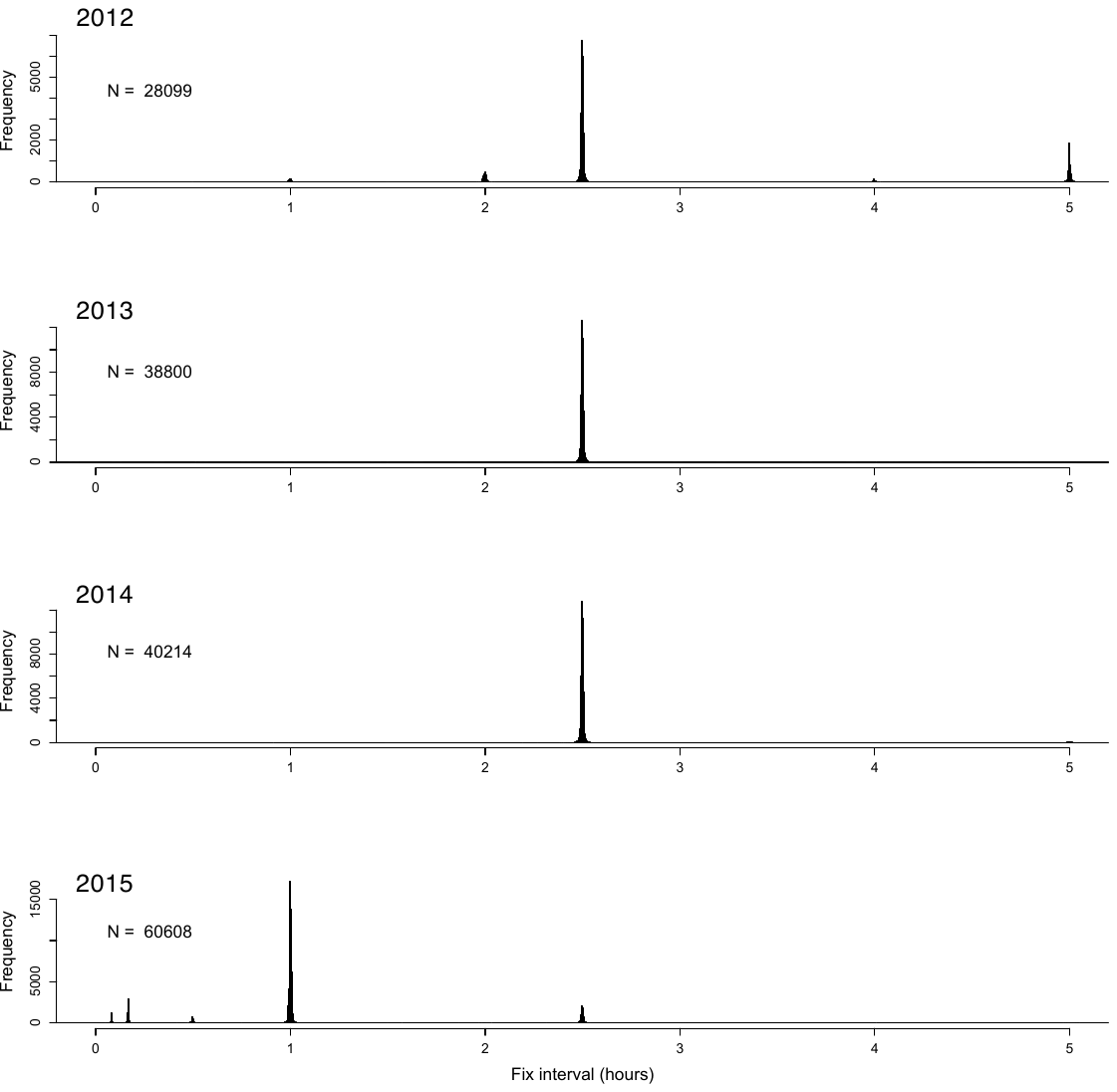

**Figure S1.** Distribution of elk GPS fix intervals for the four winters considered in this study.
