## Supplementary Materials for "Weak spatiotemporal response of prey to predation risk in a freely interacting system"

**Table S1.** Wolf pack sizes used to estimate the joint wolf utilisation distribution.

| Winter | Pack name | Size^*^ |
| --- | --- | --- |
| 2000 | Druid Peak | 27 |
|  | Leopold | 13 |
| 2001 | Leopold | 14 |
| 2002 | Druid Peak | 11 |
|  | Geode Creek | 9 |
|  | Swan Lake | 16 |
| 2003 | Druid Peak | 17 |
|  | Geode Creek | 7 |
|  | Slough Creek | 15 |
| 2004 | Geode Creek | 12 |
|  | Leopold | 23 |
|  | Loner | 1 |
| 2005 | Agate Creek | 8 |
|  | Hellroaring | 7 |
|  | Leopold | 14 |
|  | Slough Creek | 15 |
| 2006 | Agate Creek | 13 |
|  | Druid Peak | 12 |
|  | Leopold | 19 |
| 2007 | Leopold | 16 |
|  | Oxbow Creek | 16 |
| 2008 | Agate Creek | 4 |
|  | Druid Peak | 13 |
|  | Loner | 1 |
|  | Blacktail | 8 |
|  | Everts | 8 |
| 2009 | Blacktail | 9 |
| 2010 | Agate Creek | 8 |
|  | 642f Group | 1 |
|  | Blacktail | 14 |
|  | Lamar Canyon | 7 |
| 2011 | Agate Creek | 3 |
|  | Mollie's | 19 |
|  | Blacktail | 15 |
|  | Lamar Canyon | 11 |
|  | Steamboat | 4 |
|  | 777m Group (Blacktail Satellite) | 2 |
| 2012 | Blacktail | 4 |
|  | Junction Butte | 9 |
|  | Lamar Canyon | 11 |
|  | Steamboat | 3 |
|  | 8 Mile | 10 |
|  | 889f/890m Group | 2 |
| 2013 | Blacktail | 3 |
|  | Junction Butte | 9 |
|  | 755m/889f Group | 2 |
|  | 8 Mile | 18 |
|  | 911m Group | 1 |
|  | Prospect Peak | 2 |
| 2014 | Mollie's | 12 |
|  | Junction Butte | 8 |
|  | Lamar Canyon | 8 |
|  | 8 Mile | 9 |
|  | 911m Group | 2 |
|  | 967m Group | 2 |
|  | Prospect Peak | 14 |
| 2015 | Mollie's | 16 |
|  | Junction Butte | 14 |
|  | Lamar Canyon | 9 |
|  | 8 Mile | 13 |
|  | Prospect Peak | 13 |

^*^ Including adults and pups
