## Supplementary Materials for "Weak spatiotemporal response of prey to predation risk in a freely interacting system"

**Table S2.** Details of individual elk trajectories and observed and expected spatial overlap with the three predation risk layers during the four winters considered. Expected values were obtained from a null model formulation that randomised elk trajectories across the northern range, and thus did not account for philopatric behaviour. Expected distribution variances are multiplied by 100.

| Winter | Elk ID | # relocations | # tracking days | Mean # hours  between relocations | Wolf space use intensity | | | | Elk kill site density | | | | Vegetation openness | | | |
| --- | --- | --- | --- | --- | --- | --- | --- | --- | --- | --- | --- | --- | --- | --- | --- | --- |
|  |  |  |  |  | Obs. | Exp. | Var. | *P* | Obs. | Exp. | Var. | *P* | Obs. | Exp. | Var. | *P* |
| 2012 | 1201 | 1367 | 176 | 3.091 | 0.143 | 0.066 | 0.120 | 1.000 | 0.190 | 0.119 | 0.605 | 0.782 | 0.028 | 0.024 | 0.120 | 0.792 |
|  | 1203 | 1279 | 181 | 3.396 | 0.065 | 0.050 | 0.090 | 0.663 | 0.080 | 0.095 | 0.431 | 0.436 | 0.020 | 0.018 | 0.090 | 0.782 |
|  | 1204 | 1301 | 181 | 3.339 | 0.114 | 0.055 | 0.102 | 0.990 | 0.137 | 0.103 | 0.424 | 0.693 | 0.023 | 0.019 | 0.102 | 0.950 |
|  | 1205 | 1265 | 181 | 3.434 | 0.131 | 0.071 | 0.124 | 0.970 | 0.185 | 0.129 | 0.532 | 0.772 | 0.027 | 0.023 | 0.124 | 0.861 |
|  | 1206 | 1302 | 181 | 3.336 | 0.064 | 0.030 | 0.047 | 0.931 | 0.100 | 0.057 | 0.211 | 0.772 | 0.012 | 0.011 | 0.047 | 0.683 |
|  | 1207 | 1444 | 181 | 3.009 | 0.081 | 0.075 | 0.078 | 0.545 | 0.251 | 0.180 | 0.843 | 0.713 | 0.029 | 0.027 | 0.078 | 0.693 |
|  | 1208 | 1133 | 147 | 3.106 | 0.053 | 0.037 | 0.041 | 0.762 | 0.177 | 0.083 | 0.296 | 0.941 | 0.012 | 0.013 | 0.041 | 0.347 |
|  | 1209 | 1394 | 181 | 3.117 | 0.070 | 0.054 | 0.065 | 0.713 | 0.237 | 0.133 | 0.589 | 0.931 | 0.021 | 0.020 | 0.065 | 0.584 |
|  | 1210 | 1356 | 181 | 3.204 | 0.093 | 0.055 | 0.099 | 0.881 | 0.175 | 0.109 | 0.433 | 0.842 | 0.019 | 0.019 | 0.099 | 0.584 |
|  | 1211 | 1073 | 170 | 3.804 | 0.136 | 0.079 | 0.131 | 0.921 | 0.189 | 0.140 | 0.606 | 0.683 | 0.029 | 0.028 | 0.131 | 0.634 |
|  | 1212 | 1445 | 181 | 3.007 | 0.054 | 0.058 | 0.057 | 0.376 | 0.190 | 0.204 | 1.004 | 0.426 | 0.023 | 0.022 | 0.057 | 0.644 |
|  | 1214 | 1353 | 181 | 3.212 | 0.097 | 0.069 | 0.088 | 0.832 | 0.309 | 0.186 | 0.863 | 0.891 | 0.027 | 0.025 | 0.088 | 0.752 |
|  | 1215 | 1466 | 181 | 2.964 | 0.142 | 0.077 | 0.160 | 0.941 | 0.209 | 0.124 | 0.564 | 0.822 | 0.028 | 0.026 | 0.160 | 0.752 |
| 2013 | 1204 | 1724 | 181 | 2.518 | 0.105 | 0.051 | 0.082 | 0.990 | 0.156 | 0.097 | 0.375 | 0.792 | 0.019 | 0.018 | 0.001 | 0.554 |
|  | 1205 | 1736 | 181 | 2.501 | 0.107 | 0.061 | 0.104 | 0.950 | 0.149 | 0.110 | 0.408 | 0.644 | 0.023 | 0.019 | 0.001 | 0.911 |
|  | 1207 | 1736 | 181 | 2.501 | 0.085 | 0.085 | 0.157 | 0.455 | 0.172 | 0.185 | 1.133 | 0.475 | 0.037 | 0.031 | 0.001 | 0.970 |
|  | 1208 | 1312 | 138 | 2.524 | 0.028 | 0.017 | 0.019 | 0.792 | 0.110 | 0.032 | 0.086 | 0.990 | 0.006 | 0.008 | 0.000 | 0.267 |
|  | 1210 | 1585 | 170 | 2.575 | 0.119 | 0.062 | 0.099 | 0.980 | 0.234 | 0.115 | 0.465 | 0.980 | 0.022 | 0.020 | 0.001 | 0.792 |
|  | 1212 | 1710 | 180 | 2.532 | 0.035 | 0.050 | 0.033 | 0.248 | 0.135 | 0.164 | 0.366 | 0.356 | 0.019 | 0.022 | 0.001 | 0.168 |
|  | 1214 | 1683 | 181 | 2.580 | 0.064 | 0.040 | 0.048 | 0.842 | 0.131 | 0.072 | 0.226 | 0.861 | 0.015 | 0.013 | 0.001 | 0.772 |
|  | 1215 | 1661 | 173 | 2.505 | 0.120 | 0.063 | 0.136 | 0.941 | 0.200 | 0.103 | 0.464 | 0.881 | 0.023 | 0.022 | 0.001 | 0.673 |
|  | 1311 | 1685 | 170 | 2.418 | 0.035 | 0.027 | 0.038 | 0.693 | 0.128 | 0.050 | 0.146 | 0.980 | 0.010 | 0.010 | 0.000 | 0.535 |
|  | 1312 | 1688 | 176 | 2.502 | 0.042 | 0.019 | 0.015 | 0.950 | 0.091 | 0.047 | 0.105 | 0.891 | 0.009 | 0.008 | 0.000 | 0.772 |
|  | 1313 | 1693 | 181 | 2.564 | 0.005 | 0.015 | 0.019 | 0.307 | 0.041 | 0.023 | 0.055 | 0.782 | 0.007 | 0.006 | 0.000 | 0.703 |
|  | 1314 | 1724 | 181 | 2.518 | 0.082 | 0.052 | 0.082 | 0.832 | 0.182 | 0.113 | 0.583 | 0.822 | 0.020 | 0.020 | 0.001 | 0.574 |
|  | 1315 | 1681 | 181 | 2.583 | 0.072 | 0.050 | 0.065 | 0.772 | 0.234 | 0.101 | 0.357 | 0.990 | 0.019 | 0.017 | 0.001 | 0.743 |
|  | 1317 | 1733 | 181 | 2.505 | 0.079 | 0.054 | 0.066 | 0.842 | 0.188 | 0.110 | 0.513 | 0.802 | 0.020 | 0.019 | 0.001 | 0.545 |
|  | 1318 | 1736 | 181 | 2.501 | 0.016 | 0.024 | 0.040 | 0.416 | 0.010 | 0.045 | 0.142 | 0.327 | 0.013 | 0.010 | 0.000 | 0.921 |
|  | 1319 | 1736 | 181 | 2.501 | 0.027 | 0.036 | 0.053 | 0.386 | 0.083 | 0.061 | 0.201 | 0.693 | 0.015 | 0.013 | 0.001 | 0.743 |
|  | 1320 | 1783 | 181 | 2.435 | 0.089 | 0.044 | 0.074 | 0.931 | 0.180 | 0.083 | 0.390 | 0.911 | 0.019 | 0.016 | 0.001 | 0.802 |
|  | 1401 | 1335 | 139 | 2.499 | 0.006 | 0.029 | 0.034 | 0.168 | 0.036 | 0.053 | 0.135 | 0.376 | 0.014 | 0.010 | 0.000 | 0.970 |
|  | 1402 | 1335 | 139 | 2.499 | 0.032 | 0.019 | 0.025 | 0.802 | 0.030 | 0.034 | 0.111 | 0.554 | 0.008 | 0.007 | 0.000 | 0.455 |
|  | 1404 | 1335 | 139 | 2.499 | 0.055 | 0.043 | 0.059 | 0.624 | 0.086 | 0.085 | 0.320 | 0.535 | 0.018 | 0.014 | 0.001 | 0.931 |
|  | 1409 | 1307 | 139 | 2.553 | 0.033 | 0.024 | 0.033 | 0.743 | 0.033 | 0.038 | 0.097 | 0.515 | 0.008 | 0.008 | 0.000 | 0.594 |
|  | 1410 | 1335 | 139 | 2.499 | 0.023 | 0.040 | 0.049 | 0.267 | 0.046 | 0.081 | 0.297 | 0.307 | 0.020 | 0.014 | 0.001 | 1.000 |
| 2014 | 1204 | 1736 | 181 | 2.501 | 0.120 | 0.053 | 0.071 | 1.000 | 0.144 | 0.099 | 0.395 | 0.733 | 0.022 | 0.018 | 0.001 | 0.881 |
|  | 1207 | 1305 | 136 | 2.500 | 0.059 | 0.052 | 0.077 | 0.545 | 0.151 | 0.117 | 0.497 | 0.644 | 0.021 | 0.019 | 0.001 | 0.752 |
|  | 1208 | 1550 | 161 | 2.501 | 0.073 | 0.038 | 0.054 | 0.941 | 0.151 | 0.083 | 0.311 | 0.871 | 0.015 | 0.015 | 0.001 | 0.416 |
|  | 1210 | 1658 | 181 | 2.617 | 0.120 | 0.070 | 0.151 | 0.901 | 0.215 | 0.122 | 0.475 | 0.891 | 0.024 | 0.024 | 0.001 | 0.574 |
|  | 1212 | 1632 | 176 | 2.593 | 0.057 | 0.054 | 0.046 | 0.564 | 0.153 | 0.173 | 0.662 | 0.426 | 0.024 | 0.024 | 0.002 | 0.396 |
|  | 1214 | 1637 | 176 | 2.582 | 0.086 | 0.059 | 0.047 | 0.911 | 0.264 | 0.160 | 0.508 | 0.911 | 0.025 | 0.023 | 0.001 | 0.743 |
|  | 1215 | 1667 | 180 | 2.598 | 0.119 | 0.065 | 0.175 | 0.901 | 0.172 | 0.119 | 0.562 | 0.673 | 0.022 | 0.021 | 0.001 | 0.604 |
|  | 1311 | 1544 | 162 | 2.520 | 0.067 | 0.048 | 0.050 | 0.802 | 0.170 | 0.106 | 0.363 | 0.792 | 0.019 | 0.017 | 0.001 | 0.802 |
|  | 1313 | 1401 | 146 | 2.500 | 0.004 | 0.014 | 0.011 | 0.238 | 0.040 | 0.021 | 0.038 | 0.812 | 0.006 | 0.005 | 0.000 | 0.703 |
|  | 1318 | 1737 | 181 | 2.499 | 0.015 | 0.024 | 0.034 | 0.366 | 0.010 | 0.048 | 0.169 | 0.287 | 0.012 | 0.009 | 0.000 | 0.911 |
|  | 1319 | 1737 | 181 | 2.499 | 0.031 | 0.046 | 0.081 | 0.356 | 0.085 | 0.085 | 0.378 | 0.584 | 0.021 | 0.018 | 0.001 | 0.822 |
|  | 1320 | 1734 | 181 | 2.504 | 0.169 | 0.092 | 0.167 | 0.990 | 0.361 | 0.173 | 0.886 | 0.980 | 0.034 | 0.031 | 0.002 | 0.743 |
|  | 1401 | 1737 | 181 | 2.499 | 0.010 | 0.023 | 0.030 | 0.337 | 0.057 | 0.042 | 0.121 | 0.673 | 0.013 | 0.008 | 0.000 | 1.000 |
|  | 1402 | 1737 | 181 | 2.499 | 0.035 | 0.023 | 0.026 | 0.782 | 0.042 | 0.044 | 0.137 | 0.554 | 0.009 | 0.009 | 0.000 | 0.634 |
|  | 1404 | 1690 | 176 | 2.504 | 0.101 | 0.078 | 0.136 | 0.673 | 0.246 | 0.181 | 1.328 | 0.634 | 0.030 | 0.030 | 0.001 | 0.525 |
|  | 1409 | 1672 | 174 | 2.499 | 0.075 | 0.041 | 0.052 | 0.941 | 0.127 | 0.097 | 0.427 | 0.683 | 0.018 | 0.017 | 0.001 | 0.584 |
|  | 1410 | 1709 | 181 | 2.540 | 0.042 | 0.058 | 0.108 | 0.347 | 0.079 | 0.098 | 0.371 | 0.396 | 0.029 | 0.022 | 0.001 | 0.960 |
|  | 1516 | 1295 | 135 | 2.501 | 0.050 | 0.042 | 0.041 | 0.713 | 0.109 | 0.086 | 0.219 | 0.663 | 0.017 | 0.014 | 0.001 | 0.931 |
|  | 1517 | 1296 | 135 | 2.499 | 0.052 | 0.044 | 0.053 | 0.673 | 0.137 | 0.085 | 0.282 | 0.802 | 0.017 | 0.015 | 0.001 | 0.832 |
|  | 1518 | 1275 | 134 | 2.521 | 0.032 | 0.027 | 0.038 | 0.634 | 0.127 | 0.050 | 0.137 | 0.950 | 0.008 | 0.010 | 0.000 | 0.238 |
|  | 1519 | 1296 | 135 | 2.499 | 0.087 | 0.077 | 0.022 | 0.772 | 0.289 | 0.236 | 0.401 | 0.792 | 0.037 | 0.031 | 0.001 | 0.990 |
|  | 1520 | 1296 | 135 | 2.499 | 0.084 | 0.058 | 0.042 | 0.960 | 0.207 | 0.171 | 0.677 | 0.644 | 0.025 | 0.022 | 0.001 | 0.901 |
| 2015 | 1208 | 4032 | 172 | 1.025 | 0.043 | 0.024 | 0.029 | 0.832 | 0.081 | 0.044 | 0.129 | 0.822 | 0.007 | 0.009 | 0.000 | 0.277 |
|  | 1210 | 3912 | 176 | 1.078 | 0.094 | 0.042 | 0.057 | 0.980 | 0.160 | 0.079 | 0.251 | 0.960 | 0.018 | 0.015 | 0.001 | 0.891 |
|  | 1311 | 3836 | 174 | 1.088 | 0.022 | 0.015 | 0.016 | 0.703 | 0.060 | 0.027 | 0.057 | 0.891 | 0.007 | 0.005 | 0.000 | 0.743 |
|  | 1317 | 4073 | 173 | 1.020 | 0.067 | 0.057 | 0.039 | 0.713 | 0.184 | 0.162 | 0.350 | 0.663 | 0.021 | 0.019 | 0.001 | 0.772 |
|  | 1318 | 4244 | 182 | 1.029 | 0.011 | 0.016 | 0.018 | 0.436 | 0.007 | 0.024 | 0.059 | 0.386 | 0.009 | 0.006 | 0.000 | 0.921 |
|  | 1320 | 4154 | 182 | 1.051 | 0.103 | 0.054 | 0.094 | 0.970 | 0.201 | 0.108 | 0.556 | 0.842 | 0.023 | 0.020 | 0.001 | 0.871 |
|  | 1404 | 4165 | 176 | 1.013 | 0.025 | 0.044 | 0.040 | 0.218 | 0.078 | 0.118 | 0.382 | 0.327 | 0.020 | 0.016 | 0.001 | 0.941 |
|  | 1410 | 4039 | 182 | 1.081 | 0.021 | 0.036 | 0.055 | 0.317 | 0.055 | 0.075 | 0.291 | 0.396 | 0.021 | 0.014 | 0.001 | 1.000 |
|  | 1516 | 4109 | 182 | 1.062 | 0.047 | 0.030 | 0.028 | 0.832 | 0.119 | 0.051 | 0.112 | 0.970 | 0.013 | 0.011 | 0.001 | 0.762 |
|  | 1517 | 4127 | 182 | 1.058 | 0.074 | 0.052 | 0.056 | 0.891 | 0.223 | 0.118 | 0.422 | 0.931 | 0.021 | 0.017 | 0.001 | 0.941 |
|  | 1518 | 4006 | 179 | 1.074 | 0.046 | 0.035 | 0.044 | 0.634 | 0.139 | 0.070 | 0.259 | 0.891 | 0.014 | 0.012 | 0.001 | 0.782 |
|  | 1519 | 4132 | 182 | 1.056 | 0.082 | 0.073 | 0.017 | 0.743 | 0.215 | 0.208 | 0.095 | 0.574 | 0.032 | 0.026 | 0.001 | 0.960 |
