## Supplementary Materials for "Weak spatiotemporal response of prey to predation risk in a freely interacting system"

Table S4. Details of individual elk trajectories and observed and expected encounter rates during the six winter periods considered. Expected values were obtained from a null model formulation that randomised elk trajectories within the same bounding box as the original trajectory, thus accounting for philopatric behaviour.

| Winter period | Elk ID | # relocations | Mean # hours  between relocations | # encounters | Encounter rate per 100 fixes | # days per encounter | Null model | | |
| --- | --- | --- | --- | --- | --- | --- | --- | --- | --- |
|  |  |  |  |  |  |  | Exp. | Var. | *P* |
| LW 2013 | 1201 | 307 | 2.497 | 6 | 1.954 | 5.1 | 0.583 | 0.620 | 0.921 |
|  | 1202 | 307 | 2.497 | 0 | 0.000 | - | 0.000 | 0.000 | 1.000 |
|  | 1203 | 306 | 2.505 | 7 | 2.288 | 4.4 | 1.461 | 1.166 | 0.871 |
|  | 1204 | 306 | 2.505 | 0 | 0.000 | - | 1.216 | 4.814 | 0.337 |
|  | 1205 | 270 | 2.827 | 0 | 0.000 | - | 1.041 | 1.584 | 0.347 |
|  | 1206 | 306 | 2.505 | 3 | 0.980 | 10.2 | 0.967 | 1.426 | 0.752 |
|  | 1207 | 307 | 2.497 | 1 | 0.326 | 30.7 | 2.580 | 16.367 | 0.426 |
|  | 1208 | 307 | 2.497 | 0 | 0.000 | - | 1.397 | 3.602 | 0.416 |
|  | 1209 | 307 | 2.497 | 11 | 3.583 | 2.8 | 1.023 | 1.492 | 0.950 |
|  | 1210 | 288 | 2.662 | 0 | 0.000 | - | 1.222 | 3.165 | 0.327 |
|  | 1211 | 306 | 2.507 | 2 | 0.654 | 15.3 | 1.539 | 1.036 | 0.248 |
|  | 1212 | 302 | 2.538 | 0 | 0.000 | - | 0.026 | 0.015 | 0.950 |
|  | 1214 | 306 | 2.505 | 17 | 5.556 | 1.8 | 0.706 | 1.551 | 0.990 |
|  | 1215 | 307 | 2.497 | 2 | 0.651 | 15.4 | 2.541 | 6.964 | 0.228 |
|  | 1311 | 306 | 2.505 | 5 | 1.634 | 6.1 | 0.775 | 1.676 | 0.881 |
|  | 1312 | 307 | 2.505 | 0 | 0.000 | - | 0.541 | 0.678 | 0.505 |
|  | 1313 | 307 | 2.505 | 3 | 0.977 | 10.2 | 1.016 | 0.674 | 0.604 |
|  | 1314 | 307 | 2.505 | 7 | 2.280 | 4.4 | 2.788 | 11.080 | 0.653 |
|  | 1315 | 307 | 2.505 | 7 | 2.280 | 4.4 | 1.342 | 4.095 | 0.772 |
|  | 1316 | 307 | 2.505 | 4 | 1.303 | 7.7 | 3.173 | 13.357 | 0.495 |
|  | 1317 | 307 | 2.505 | 0 | 0.000 | - | 0.088 | 0.085 | 0.861 |
|  | 1318 | 307 | 2.505 | 15 | 4.886 | 2.0 | 4.300 | 16.775 | 0.663 |
|  | 1319 | 307 | 2.498 | 2 | 0.651 | 15.4 | 0.687 | 1.057 | 0.713 |
|  | 1320 | 306 | 2.513 | 16 | 5.229 | 1.9 | 1.307 | 3.102 | 0.960 |
| EW 2013 | 1202 | 308 | 2.500 | 0 | 0.000 | - | 0.081 | 0.112 | 0.881 |
|  | 1204 | 308 | 2.500 | 7 | 2.273 | 4.4 | 2.737 | 4.217 | 0.515 |
|  | 1205 | 308 | 2.500 | 10 | 3.247 | 3.1 | 1.899 | 3.470 | 0.822 |
|  | 1207 | 308 | 2.500 | 1 | 0.325 | 30.8 | 2.078 | 6.842 | 0.188 |
|  | 1210 | 297 | 2.593 | 6 | 2.020 | 5.0 | 2.226 | 2.350 | 0.535 |
|  | 1212 | 308 | 2.500 | 13 | 4.221 | 2.4 | 1.786 | 4.416 | 0.901 |
|  | 1214 | 285 | 2.703 | 2 | 0.702 | 14.3 | 1.056 | 0.650 | 0.554 |
|  | 1215 | 308 | 2.500 | 8 | 2.597 | 3.9 | 2.494 | 2.525 | 0.604 |
|  | 1311 | 364 | 2.114 | 1 | 0.275 | 36.4 | 1.008 | 1.211 | 0.396 |
|  | 1312 | 308 | 2.500 | 7 | 2.273 | 4.4 | 2.302 | 2.671 | 0.614 |
|  | 1313 | 298 | 2.584 | 3 | 1.007 | 9.9 | 0.540 | 0.514 | 0.881 |
|  | 1314 | 308 | 2.500 | 2 | 0.649 | 15.4 | 1.912 | 2.941 | 0.327 |
|  | 1315 | 302 | 2.550 | 3 | 0.993 | 10.1 | 1.636 | 2.290 | 0.525 |
|  | 1316 | 308 | 2.500 | 2 | 0.649 | 15.4 | 1.432 | 1.312 | 0.356 |
|  | 1317 | 308 | 2.500 | 3 | 0.974 | 10.3 | 1.390 | 1.844 | 0.535 |
|  | 1318 | 308 | 2.500 | 3 | 0.974 | 10.3 | 0.244 | 0.241 | 0.980 |
|  | 1319 | 308 | 2.500 | 0 | 0.000 | - | 1.386 | 2.762 | 0.248 |
|  | 1320 | 371 | 2.074 | 4 | 1.078 | 9.3 | 1.633 | 2.259 | 0.495 |
| LW 2014 | 1202 | 306 | 2.505 | 0 | 0.000 | - | 0.000 | 0.000 | 1.000 |
|  | 1204 | 306 | 2.505 | 0 | 0.000 | - | 0.961 | 1.900 | 0.465 |
|  | 1205 | 306 | 2.505 | 1 | 0.327 | 30.6 | 0.559 | 0.866 | 0.683 |
|  | 1207 | 306 | 2.505 | 0 | 0.000 | - | 0.219 | 0.384 | 0.802 |
|  | 1208 | 306 | 2.505 | 5 | 11.111 | 6.1 | 4.101 | 11.408 | 0.970 |
|  | 1210 | 287 | 2.671 | 1 | 0.348 | 28.7 | 2.251 | 3.920 | 0.198 |
|  | 1212 | 306 | 2.505 | 1 | 0.327 | 30.6 | 0.399 | 0.787 | 0.822 |
|  | 1214 | 306 | 2.505 | 4 | 1.307 | 7.7 | 2.301 | 5.072 | 0.455 |
|  | 1215 | 305 | 2.513 | 0 | 0.000 | - | 1.023 | 1.247 | 0.356 |
|  | 1311 | 306 | 2.505 | 7 | 2.288 | 4.4 | 2.958 | 8.555 | 0.574 |
|  | 1312 | 305 | 2.513 | 2 | 0.656 | 15.3 | 2.020 | 4.143 | 0.465 |
|  | 1313 | 306 | 2.505 | 4 | 1.307 | 7.7 | 0.797 | 0.676 | 0.851 |
|  | 1314 | 306 | 2.505 | 1 | 0.327 | 30.6 | 1.637 | 2.288 | 0.386 |
|  | 1315 | 299 | 2.564 | 5 | 1.672 | 6.0 | 2.749 | 8.961 | 0.515 |
|  | 1317 | 306 | 2.505 | 0 | 0.000 | - | 2.941 | 5.983 | 0.059 |
|  | 1318 | 306 | 2.505 | 4 | 1.307 | 7.7 | 0.761 | 1.185 | 0.802 |
|  | 1319 | 306 | 2.505 | 3 | 0.980 | 10.2 | 0.271 | 0.138 | 0.970 |
|  | 1320 | 306 | 2.505 | 0 | 0.000 | - | 3.458 | 10.164 | 0.119 |
|  | 1401 | 307 | 2.497 | 0 | 0.000 | - | 0.059 | 0.059 | 0.931 |
|  | 1402 | 307 | 2.497 | 12 | 3.909 | 2.6 | 1.788 | 1.371 | 0.980 |
|  | 1404 | 307 | 2.497 | 0 | 0.000 | - | 0.779 | 1.155 | 0.416 |
|  | 1409 | 300 | 2.555 | 8 | 2.667 | 3.8 | 1.743 | 3.329 | 0.713 |
|  | 1410 | 307 | 2.497 | 0 | 0.000 | - | 0.404 | 0.603 | 0.644 |
| EW 2014 | 1202 | 304 | 2.533 | 1 | 0.329 | 30.4 | 0.490 | 0.585 | 0.683 |
|  | 1204 | 308 | 2.500 | 7 | 2.273 | 4.4 | 1.406 | 1.597 | 0.792 |
|  | 1207 | 308 | 2.500 | 6 | 1.948 | 5.1 | 1.698 | 2.349 | 0.673 |
|  | 1210 | 291 | 2.647 | 0 | 0.000 | - | 1.261 | 1.234 | 0.149 |
|  | 1212 | 289 | 2.665 | 1 | 0.346 | 28.9 | 1.215 | 1.852 | 0.426 |
|  | 1214 | 284 | 2.703 | 16 | 5.634 | 1.8 | 2.218 | 3.773 | 0.950 |
|  | 1215 | 280 | 2.751 | 1 | 0.357 | 28.0 | 1.439 | 1.442 | 0.307 |
|  | 1313 | 308 | 2.500 | 0 | 0.000 | - | 2.159 | 3.640 | 0.089 |
|  | 1314 | 308 | 2.500 | 0 | 0.000 | - | 0.630 | 0.549 | 0.327 |
|  | 1315 | 308 | 2.500 | 14 | 4.545 | 2.2 | 2.010 | 2.774 | 0.931 |
|  | 1318 | 308 | 2.500 | 19 | 6.169 | 1.6 | 2.688 | 5.173 | 0.931 |
|  | 1319 | 308 | 2.500 | 8 | 2.597 | 3.9 | 1.338 | 2.073 | 0.822 |
|  | 1320 | 308 | 2.500 | 5 | 1.623 | 6.2 | 1.490 | 2.505 | 0.663 |
|  | 1401 | 308 | 2.500 | 1 | 0.325 | 30.8 | 1.568 | 1.619 | 0.228 |
|  | 1402 | 308 | 2.500 | 5 | 1.623 | 6.2 | 2.039 | 3.146 | 0.554 |
|  | 1404 | 308 | 2.500 | 1 | 0.325 | 30.8 | 1.575 | 1.474 | 0.208 |
|  | 1409 | 308 | 2.500 | 5 | 1.623 | 6.2 | 1.623 | 1.944 | 0.624 |
|  | 1410 | 308 | 2.500 | 2 | 0.649 | 15.4 | 1.380 | 1.830 | 0.416 |
| LW 2015 | 1202 | 300 | 2.555 | 0 | 0.000 | - | 0.000 | 0.000 | 1.000 |
|  | 1204 | 307 | 2.497 | 5 | 1.629 | 6.1 | 1.798 | 1.675 | 0.564 |
|  | 1208 | 307 | 2.497 | 0 | 0.000 | - | 0.954 | 1.355 | 0.267 |
|  | 1210 | 289 | 2.653 | 4 | 1.384 | 7.2 | 0.886 | 1.007 | 0.802 |
|  | 1212 | 301 | 2.547 | 0 | 0.000 | - | 0.000 | 0.000 | 1.000 |
|  | 1214 | 307 | 2.497 | 2 | 0.651 | 15.4 | 1.752 | 3.056 | 0.386 |
|  | 1215 | 295 | 2.599 | 5 | 1.695 | 5.9 | 1.390 | 1.315 | 0.703 |
|  | 1311 | 295 | 2.599 | 6 | 2.034 | 4.9 | 2.766 | 4.271 | 0.485 |
|  | 1314 | 307 | 2.497 | 0 | 0.000 | - | 0.000 | 0.000 | 1.000 |
|  | 1318 | 307 | 2.497 | 14 | 4.560 | 2.2 | 4.221 | 10.612 | 0.564 |
|  | 1319 | 307 | 2.497 | 0 | 0.000 | - | 0.446 | 0.679 | 0.604 |
|  | 1320 | 306 | 2.505 | 9 | 2.941 | 3.4 | 3.458 | 7.750 | 0.594 |
|  | 1401 | 307 | 2.497 | 3 | 0.977 | 10.2 | 0.625 | 0.790 | 0.802 |
|  | 1402 | 307 | 2.497 | 0 | 0.000 | - | 0.726 | 1.099 | 0.545 |
|  | 1404 | 307 | 2.497 | 0 | 0.000 | - | 0.518 | 0.738 | 0.574 |
|  | 1409 | 307 | 2.497 | 0 | 0.000 | - | 0.580 | 0.766 | 0.535 |
|  | 1410 | 294 | 2.607 | 1 | 0.340 | 29.4 | 0.704 | 0.709 | 0.564 |
|  | 1516 | 307 | 2.497 | 10 | 3.257 | 3.1 | 3.068 | 5.816 | 0.604 |
|  | 1517 | 307 | 2.497 | 13 | 4.235 | 2.4 | 2.713 | 4.444 | 0.772 |
|  | 1518 | 296 | 2.590 | 3 | 1.014 | 9.9 | 1.118 | 1.989 | 0.713 |
|  | 1519 | 307 | 2.497 | 1 | 0.326 | 30.7 | 1.121 | 2.562 | 0.495 |
|  | 1520 | 307 | 2.497 | 0 | 0.000 | - | 0.433 | 0.707 | 0.634 |
| EW 2015 | 1208 | 298 | 2.584 | 0 | 0.000 | - | 0.151 | 0.083 | 0.673 |
|  | 1210 | 286 | 2.693 | 3 | 1.049 | 9.5 | 1.577 | 3.017 | 0.594 |
|  | 1311 | 304 | 2.533 | 0 | 0.000 | - | 0.283 | 0.127 | 0.465 |
|  | 1314 | 324 | 2.375 | 1 | 0.309 | 32.4 | 0.281 | 0.337 | 0.812 |
|  | 1317 | 314 | 2.450 | 1 | 0.318 | 31.4 | 0.742 | 0.791 | 0.554 |
|  | 1318 | 331 | 2.324 | 0 | 0.000 | - | 0.000 | 0.000 | 1.000 |
|  | 1319 | 308 | 2.500 | 3 | 0.974 | 10.3 | 1.581 | 2.333 | 0.485 |
|  | 1320 | 312 | 2.466 | 0 | 0.000 | - | 1.035 | 1.664 | 0.297 |
|  | 1401 | 324 | 2.375 | 0 | 0.000 | - | 0.164 | 0.149 | 0.792 |
|  | 1404 | 311 | 2.474 | 1 | 0.322 | 31.1 | 0.961 | 2.248 | 0.535 |
|  | 1410 | 317 | 2.427 | 0 | 0.000 | - | 0.461 | 1.064 | 0.743 |
|  | 1516 | 314 | 2.450 | 0 | 0.000 | - | 0.662 | 1.091 | 0.356 |
|  | 1517 | 332 | 2.317 | 2 | 0.602 | 16.6 | 0.527 | 0.726 | 0.802 |
|  | 1518 | 305 | 2.520 | 1 | 0.328 | 30.5 | 0.548 | 0.767 | 0.713 |
|  | 1519 | 311 | 2.474 | 3 | 0.965 | 10.4 | 1.042 | 0.813 | 0.594 |
